# A prebiotic basis for ATP as the universal energy currency

**DOI:** 10.1101/2021.10.06.463298

**Authors:** Silvana Pinna, Cäcilia Kunz, Stuart Harrison, Sean F. Jordan, John Ward, Finn Werner, Nick Lane

## Abstract

ATP is universally conserved as the principal energy currency in cells, driving metabolism through phosphorylation and condensation reactions. Such deep conservation suggests that ATP arose at an early stage of biochemical evolution. Yet purine synthesis requires six phosphorylation steps linked to ATP hydrolysis. This autocatalytic requirement for ATP to synthesize ATP implies the need for an earlier prebiotic ATP-equivalent, which could drive protometabolism before purine synthesis. Why this early phosphorylating agent was replaced, and specifically with ATP rather than other nucleotide triphosphates, remains a mystery. Here we show that the deep conservation of ATP reflects its prebiotic chemistry in relation to another universally conserved intermediate, acetyl phosphate, which bridges between thioester and phosphate metabolism by linking acetyl CoA to the substrate-level phosphorylation of ADP. We confirm earlier results showing that acetyl phosphate can phosphorylate ADP to ATP at nearly 20 % yield in water in the presence of Fe^3+^ ions. We then show that Fe^3+^ and acetyl phosphate are surprisingly favoured: a panel of other prebiotically relevant ions and minerals did not catalyze ADP phosphorylation; nor did a number of other potentially prebiotic phosphorylating agents. Only carbamoyl phosphate showed some modest phosphorylating activity. Critically, we show that acetyl phosphate does not phosphorylate other nucleotide diphosphates or free pyrophosphate in water. The phosphorylation of ADP monomers seems to be favoured by the interaction between the N6 amino group on the adenine ring with Fe^3+^ coupled to acetyl phosphate. Our findings suggest that the reason ATP is universally conserved across life is that its formation is chemically favoured in aqueous solution under mild prebiotic conditions.

## Introduction

ATP is casually referred to as the ‘universal energy currency’ of life. Why it gained this ascendency in metabolism, in place of many possible equivalents, is an abiding mystery in biology. There is nothing particularly special about the ‘high-energy’ phosphoanhydride bonds in ATP. Rather, its ability to drive phosphorylation or condensation reactions reflects the extraordinary disequilibrium between ATP and ADP – about 10 orders of magnitude in modern cells, pushed by free energy derived from respiration [1]. ATP drives intermediary metabolism through the coupling of exergonic to endergonic reactions via phosphorylation and hydrolysis, but other phosphorylating agents (including GTP and CTP) could be pushed equally far from equilibrium, and accomplish equivalent coupling. In fact, the centrality of ATP goes far beyond phosphorylation, as emphasised by the ubiquity of ATP derivatives in intermediary metabolism, including the ancient cofactors NADH, FADH and Coenzyme A (which all derive from ATP rather than AMP or adenine). ATP-coupled monomer activation also promotes the polymerisation of macromolecules, including RNA, DNA and proteins. Protein synthesis requires the activation of amino acids by adenylation (using ATP) before binding to tRNA, while the nucleotide triphosphates used for RNA and DNA synthesis are phosphorylated by ATP. So what, if anything, is special about ATP?

The most pleasing partial answer to this question is that ATP links energy metabolism with genetic information [2]. The ability to replicate RNA or DNA depends on the availability of sufficient energy to complete the task. Unlike the simple phosphorylation of intermediary metabolites, the leaving group during nucleotide polymerization is pyrophosphate (PPi) [3]. Likewise, activation of amino acids by adenylation liberates PPi as the leaving group [4–7]. The hydrolysis of PPi renders these steps exothermic, if not practically irreversible [3,8]. Only nucleotide triphosphates can release PPi while still retaining a phosphate for the sugar-phosphate backbone of RNA and DNA, or for amino-acid activation. But the fact that the canonical nucleotides can all form triphosphates, with equivalent free-energy profiles, only serves to emphasise the prominence of ATP over GTP, TTP, UTP or CTP in RNA, DNA and protein synthesis. While GTP is not uncommon in metabolic processes, including gluconeogenesis and the Krebs cycle, as well as in association with G proteins, and as the precursor of folate and pterin cofactors [9], it hardly displaces ATP from its central position in biology. Even if only one nucleotide triphosphate can be dominant, the implication of a frozen accident is not a satisfying explanation. In any case, the fact that ATP is universally conserved in the synthesis of RNA, DNA and proteins suggests it arose very early in biology, possibly even in a ‘monomer world’, before these macromolecules existed [10,11].

The mechanisms of ATP synthesis could give insight into why ATP is universally conserved. The ATP synthase is ancient and was most likely present in the last universal common ancestor of life (LUCA) [12]. But as a rotating multi-subunit nanomotor powered by the proton-motive force, the ATP synthase is clearly a product of genes and natural selection. Because LUCA had genes and molecular machines such as ribosomes, there is no inconsistency here [13]. Yet prebiotic precursors of the ATP synthase are hard to imagine [14]. This dead-end is compounded by the inference that glycolytic ATP synthesis is less deeply conserved than chemiosmotic coupling. In bacteria and archaea, many genes in both the Embden-Meyerhof-Parnas and Entner-Doudoroff pathways are not homologous, which suggests that gluconeogenesis preceded glycolysis [15,16], and that LUCA might not have had a genetically encoded glycolytic pathway. Arguably the most plausible ancestral mechanism of ATP synthesis is through the substrate-level phosphorylation of ADP to ATP by acetyl phosphate (AcP)[14], which still acts as a bridge between thioester and phosphate metabolism in bacteria and archaea [17,18]. In modern bacteria, AcP is formed by the phosphorolysis of acetyl CoA; in archaea and eukaryotes, AcP remains bound to the active site of the enzyme, but is still formed as a transient intermediate [18]. The notion that AcP played an important role at the origin of life goes back to Lipmann [19], and has been advocated by de Duve, Ferry and House, Martin and Russell, and others [17,18,20–23]. It is at least possible to imagine the substrate-level phosphorylation of ADP to ATP by AcP in a monomer world.

CoA itself is derived from ATP, but simpler thioesters, with equivalent functional chemistry to acetyl CoA, have long been linked with prebiotic chemistry and the core metabolic networks in cells [13,17,19,24–30]. Recent work suggests that thioesters such as methyl thioacetate can be synthesised under hydrothermal conditions [31]. AcP can also be made in water under ambient or mild hydrothermal conditions by phosphorolysis of thioacetate which, as a thiocarboxylic acid, is even simpler than thioesters [10]. AcP will phosphorylate various nucleotide precursors in water, including ribose to ribose-5-phosphate, and adenosine to AMP [10]. Importantly, AcP will phosphorylate ADP to ATP at 20 % yield in water in the presence of Fe^3+^ ions, suggesting that substrate-level phosphorylation could indeed take place in aqueous prebiotic conditions [32,33]. But there are also some confounding issues with AcP chemistry. Most notably, AcP acetylates amino groups, especially under alkaline conditions, which could interfere with the activation and polymerization of amino acids [10,34]. This propensity to acetylate amino acids might explain why AcP is retained in the active site of acetate kinase in archaea (and pyruvate dehydrogenase in eukaryotes) [18,35–37].

The discovery that AcP can phosphorylate ADP to ATP in the presence of Fe^3+^ was serendipitous: while studying the electrolysis of ADP in the presence of AcP, Kitani *et al*. noted a ∼20 % conversion of ADP to ATP as the iron electrode they were using in their setup corroded [32]. But the fact that substrate-level phosphorylation of ADP to ATP can be accomplished by AcP in water says nothing about whether this mechanism actually holds prebiotic relevance. We have therefore explored the phosphorylation of ADP more systematically using a range of prebiotically plausible and biologically relevant phosphorylating agents, and a panel of metal ions as possible catalysts. We find that the combination of Fe^3+^ and AcP is unique: no other metal ions or phosphorylating agents are as effective at phosphorylating ADP. Equally striking, we find that ADP is also unique: the combination of AcP and Fe^3+^ will phosphorylate ADP but not GDP, CDP, UDP or IDP, nor free pyrophosphate. We use these data and the reaction kinetics to propose a possible mechanism. Our results suggest that ATP became established as the universal energy currency in a prebiotic, monomeric world, on the basis of its unusual chemistry in water.

## Results

### Fe^3+^ is unique in promoting ADP phosphorylation by acetyl phosphate

We analysed a panel of metal ions commonly used as cofactors in metabolism, and likely available at the origin of life, to compare their effect on the phosphorylation of ADP by AcP. We first confirmed the results of Kitani *et al*. [32,33] in demonstrating that Fe^3+^ catalyses the formation of ATP by AcP at ∼15-20% yield depending on the conditions (**Fig. 1a**). We corroborated our HPLC results using MS/MS (**Fig. 1b**). Surprisingly, we found that Fe^3+^ is uniquely effective at catalysing ADP phosphorylation, at least among the large panel of metal ions we tested. FeS clusters chelated by monomeric cysteine initially seemed to produce small yields of ATP, as shown in **Fig. 1a**. However, Cys-FeS clusters are unstable and break down over hours except under strictly anoxic conditions [38]. We therefore suspected that the ATP yield actually reflected the release of Fe^3+^ into the medium. This was confirmed under more strictly anoxic conditions in an anaerobic glovebox, wherein FeS clusters failed to catalyse ATP formation (**SI Fig. 1**).

**Figure 1.**
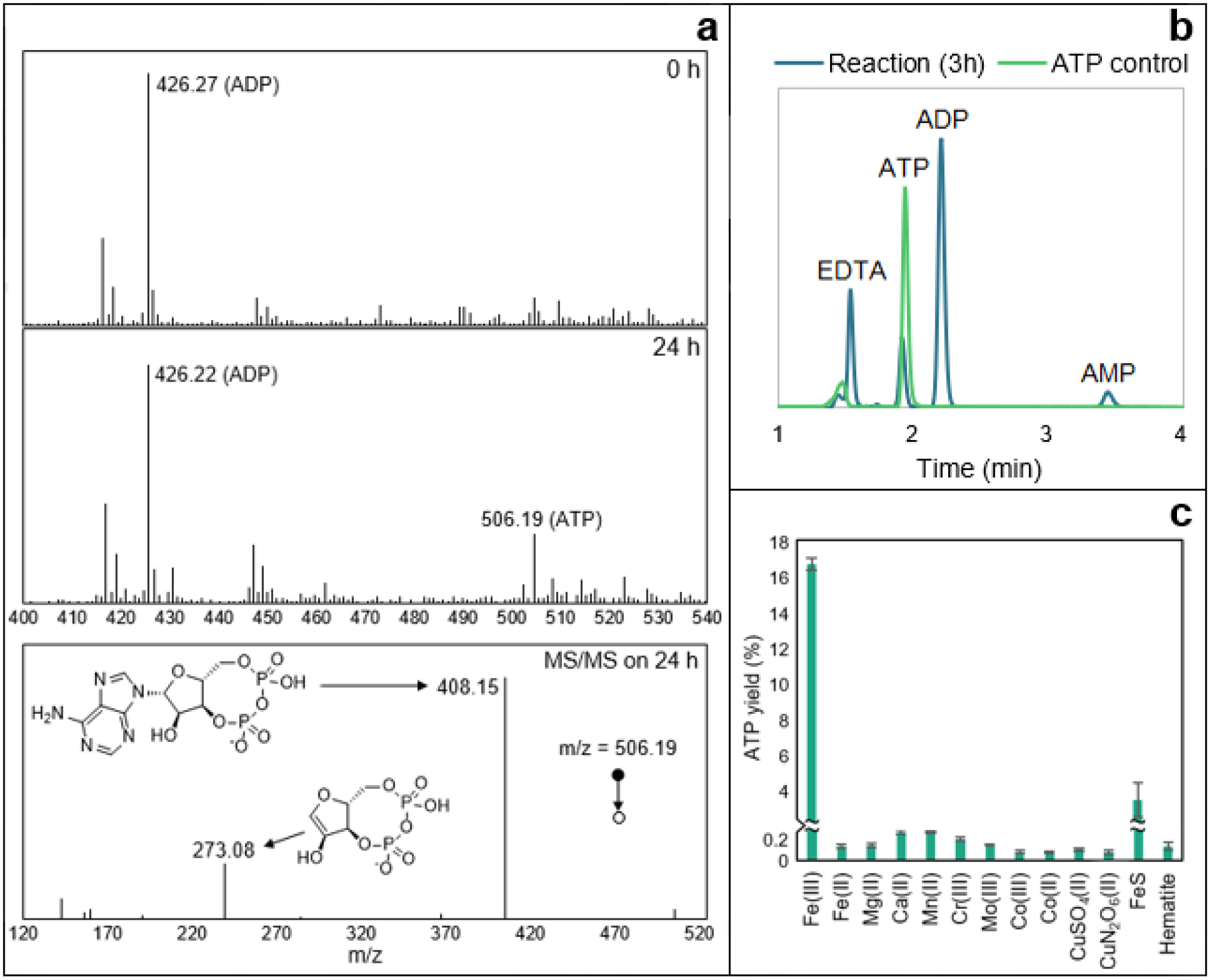
ATP synthesis with metal ion catalysts. (**a**) Mass spectrometry analysis on a reaction sample at t = 0 h (upper panel) and 24 h (middle panel). The MS/MS spectrum and proposed structures of the products of the fragmentation of the ATP mass detected at 24 h (m/z = 506.19) is shown in the lower panel and was confronted to commercial standards and public data [102]. Conditions: ADP (1 mM) + AcP (4 mM) + Fe^3+^ (500 μM) at 30°C and pH ∼5.5–6. (**b**) HPLC trace of ATP control (0.7 mM) and ATP produced by the reaction ADP (1 mM) + AcP (4 mM) + Fe^3+^ (500 μM) at 30°C and pH ∼5.5–6. (**c**) Test of reaction ADP (1 mM) + AcP (4 mM) at 30°C and pH ∼5.5–6 with Fe^3+^ (Fe_2_(SO_4_)_3_), Mg^2+^ (MgCl_2_), Ca^2+^ (CaCl_2_), Mn^2+^ (Mn(NO_3_)_2_), Cr^3+^ (Cr(NO_3_)_3_), Mo^3+^ (MoCl_3_), Co^3+^ ([Co(NH3)6]Cl3), Co^2+^ (CoCl_2_), CuSO_4_, Cu(NO_3_)_2_, FeS clusters (500 μM) and hematite (Fe_2_O_3_, 50 mg). The bars represent the ATP yield after 5h. N = 3 ±SD.

Metal ions that are commonly associated with ATP in metabolism, notably Mg^2+^ [39,40] failed to catalyse ATP formation either as free ions, or when coordinated by the monomeric amino acid aspartate, or in mineral form as brucite (**SI Fig. 2**). We had anticipated that chelated metal ions would show a stronger catalytic efficacy than free ions, as the coordination environment partially mimics the active site of enzymes, in this case acetate kinase or RNA polymerase (where glutamate or aspartate chelates Mg^2+^ at the active site). Brucite is a hydroxide mineral (Mg(OH)_2_) with a unit-cell structure that is also reminiscent of the Mg^2+^ coordination by the carboxylate of aspartate in the RNA polymerase. Surface catalysis may play an important role in prebiotic chemistry, but in this case failed to promote ATP synthesis. Mn^2+^, which has a similar activity to Mg^2+^ in acetate kinase [41] also failed to promote ATP synthesis.

### ADP phosphorylation occurs in a range of aqueous prebiotic environments

We next explored the conditions under which Fe^3+^ catalyses the phosphorylation of ADP by acetyl phosphate, specifically pH, temperature, water activity and pressure. We found that the reaction is strongly sensitive to pH, and occurs most readily under mildly acidic conditions, with an optimum pH of ∼5.5–6, the uncorrected default pH of the reaction (**Fig. 2a**). Slightly more acidic conditions (pH 4) suppressed the yield a little, but more alkaline conditions had a much stronger suppressive effect. ATP yield fell by around three quarters at pH 7, and collapsed to nearly zero at pH 9. While this sharp sensitivity to pH might seem at first sight limiting, in the Discussion we show that, on the contrary, it could be valuable in generating disequilibria, enabling ATP hydrolysis to power work.

**Figure 2.**
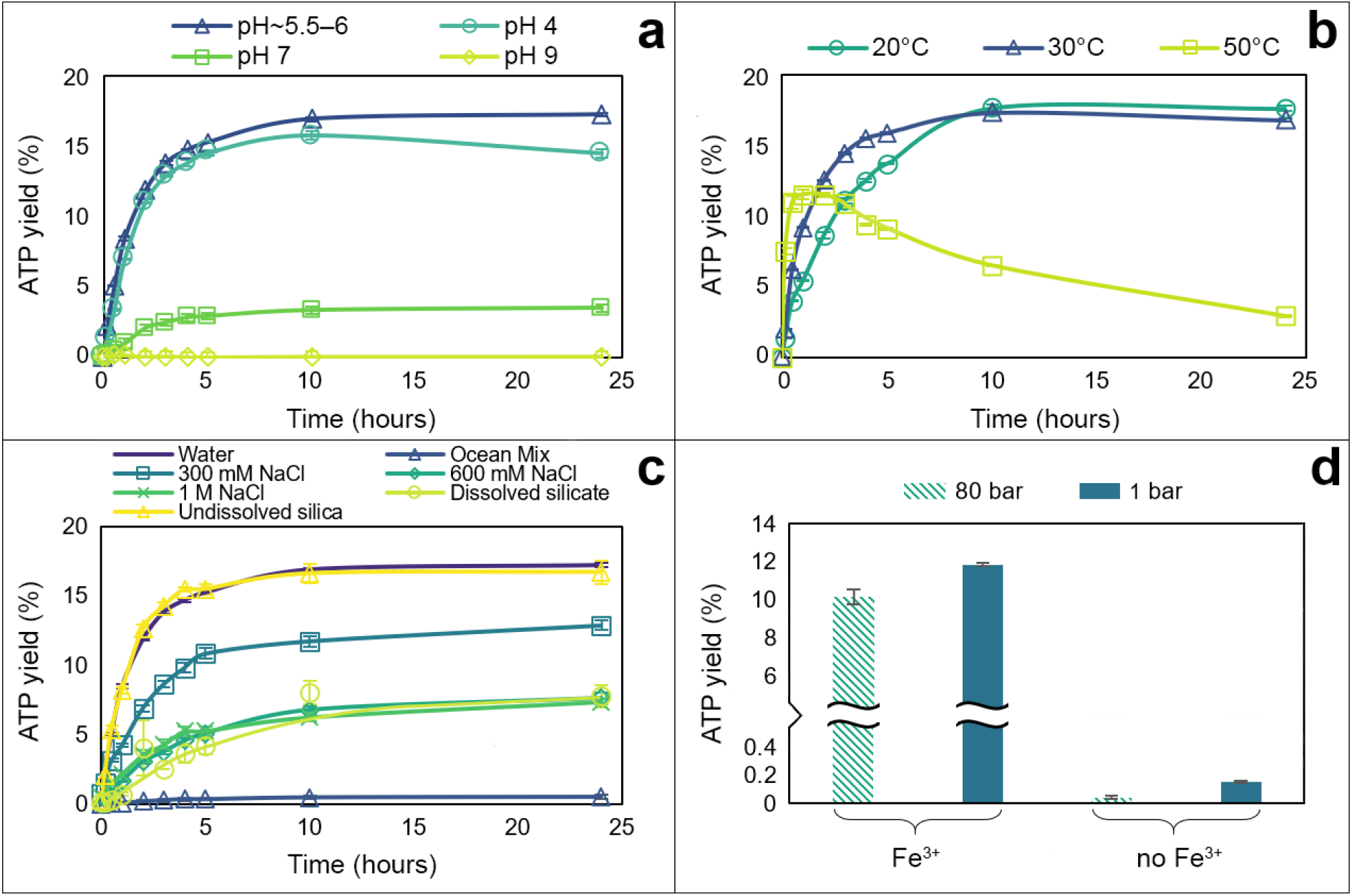
ATP synthesis by AcP and Fe^3+^ at different conditions. (**a**) Effect of pH on reaction ADP (1 mM) + AcP (4 mM) + Fe^3+^ (500 μM) at 30°C. The optimal pH of the reaction is ∼5.5–6. Rate of reaction: 0.0079 μM/s (optimal pH), 0.0074 μM/s (pH 4) and 0.0011 μM/s (pH 7). N = 3 ±SD. (**b**) Effect of temperature on reaction ADP (1 mM) + AcP (4 mM) + Fe^3+^ (500 μM), pH ∼5.5–6. Rate of reaction: 0.0066 μM/s (20°C), 0.0079 μM/s (30°C) and 0.028 μM/s (50°C). N = 3 ±SD. (**c**) Comparison of ATP yield from the reaction ADP (1 mM) + AcP (4 mM) at 30°C, pH ∼5.5–6 in water (reaction ionic strength = 3.75 mM), a modern ocean mix (600 mM NaCl, 50 mM MgCl_2_ and 10 mM CaCl_2_, reaction ionic strength = 783.75 mM), 300 mM NaCl (reaction ionic strength = 303.75 mM), modern ocean concentration of NaCl (600 mM, reaction ionic strength = 603.75 mM), 1 mM NaCl (reaction ionic strength = 1.004 M), dissolved silicate (10 mM SiO_2_, reaction ionic strength = 123.75 mM), and suspended silica in water (50 mg). N = 3 ±SD. (**d**) Comparison of ATP yield from the reaction ADP (1 mM) + AcP (4 mM) at 30°C and pH ∼5.5–6 with and without Fe^3+^ (500 µM) at 80 bar (striped yellow) and at atmospheric pressure (1 bar, solid blue). N = 2 ±SD.

ATP yield was less acutely sensitive to temperature, at least between 20 and 50 °C. Over 24 hours, the overall ATP yield reflects both synthesis and hydrolysis. We found that 30 °C optimised yield across 24 hours, by promoting synthesis within the first 4 hours while limiting hydrolysis over the subsequent 20 hours (**Fig. 2b**). The rate of synthesis was a little lower at 20 °C, but this was offset by slightly less hydrolysis over 24 hours. ATP synthesis was markedly faster at 50 °C, but so too was hydrolysis, which already lowered yields within the first 2 hours and cut them to less than a quarter of those at 30 °C after 24 hours. If ATP is to power work, as in modern cells, then hydrolysis in itself is not an issue, but rather needs to be coupled to other reactions such as the phosphorylation or condensation of substrates. Such processes also tend to take place over minutes to hours [10], meaning that temperature has a relatively trivial effect, with the yield after 2-3 hours being similar at all three temperatures studied, at around 10-15 % (**Fig. 2b**). This implies that temperature would not be a strong limiting factor on many possible prebiotic environments.

More surprisingly, ATP yield was greatest at high water activity, either in HPLC-grade water or in suspended silica (**Fig. 2c**). Adding NaCl lowered ATP yield, albeit not dramatically. Moderate NaCl concentration (300 mM, giving a total reaction ionic strength of 303.75 mM) lowered ATP yield by around a fifth. Modern ocean salinity (600 mM NaCl, reaction ionic strength 603.75 mM) and higher salinity (1 M NaCl, reaction ionic strength 1.004 M) both roughly halved the yield. This suggests that the effect of solutes does not only reflect ionic strength, which was confirmed by the addition of other solutes. Dissolved silicate (10 mM SiO_2_) also halved ATP yield, even though the ionic strength in this case was only 123.75 mM (**Fig. 2c**). Likewise, higher Mg^2+^ and Ca^2+^ concentrations (50 mM and 10 mM, respectively) as part of a modern ocean mix collapsed ATP yields to nearly zero (**Fig. 2c**), presumably because Ca^2+^ and Mg^2+^ promote ATP hydrolysis [42,43]. While this might suggest that ATP synthesis could not occur in modern oceans, Mg^2+^ and Ca^2+^ concentrations can in fact vary considerably in ocean environments (see Discussion). We show later that lower Mg^2+^ and Ca^2+^ concentrations (∼2 mM) actually promote ATP synthesis.

High pressure (80 bar) had very little effect on ATP synthesis (**Fig. 2d**). This is consistent with the work of Leibrock, Bayer, and Lüdemann (1995), who showed that high pressure promotes ATP hydrolysis, but only at pressures ≥ 300 bar. The slightly greater ATP yield at ambient pressure in our experiment may be attributable to greater evaporation in the open (non-pressurized) system. This was clearly the case in the absence of Fe^3+^, where most of the ATP detected was not produced by phosphorylation of ADP, but contamination of the ADP commercial standard via the manufacturing process, then concentrated by evaporation at ambient pressure (**SI Fig. 3**).

### Acetyl phosphate is more effective than other prebiotic phosphorylating agents

We compared AcP with a panel of eight other potentially prebiotic phosphorylating agents, including a number still used by cells today (**Table 1**).

**Table 1.** Phosphorylating agents tested.

| Name | ID | Formula | Prebiotic/biochemical prominence |
| --- | --- | --- | --- |
| Cyclic trimetaphosphate | cTMP | $\text{Na}_3\text{P}_3\text{O}_9$ | [40,45–47] |
| Pyrophosphate | PPI(V) | $\text{K}_4\text{P}_2\text{O}_7$ | [48] |
| Pyrophosphite | PPI(III) | $\text{Na}_2\text{H}_2\text{P}_2\text{O}_5$ | Has been detected in meteorites and can be generated from phosphite under hot acidic hydrothermal conditions; phosphate can be reduced to phosphite by serpentinization [49–53] |
| Phosphoenolpyruvate | PEP | $\text{KC}_3\text{H}_5\text{O}_6\text{P}$ | Has the highest phosphoryl-transfer potential found in living organisms ($\Delta G^\circ = -62$ kJ/mol) [54], and is an intermediate in gluconeogenesis and glycolysis, where its conversion to pyruvic acid by pyruvate kinase generates ATP via substrate-level phosphorylation |
| Carbamoyl phosphate | CP | $\text{Li}_2\text{CH}_2\text{NO}_5\text{P} \cdot x\text{H}_2\text{O}$ | Can be made abiotically and has a role in extant biochemistry [55] |
| Trimethyl phosphate | TMP | $(\text{CH}_3)_3\text{PO}_4$ | Has been studied for its potential role in the non-enzymatic conversion of hypoxanthine to adenine [56] |

Given the diverse reaction kinetics anticipated with these different phosphorylating agents, we carried out experiments at both at 30 °C (the optimal temperature for AcP) and 50 °C (as most phosphate donors are less labile than AcP and so might be more effective at higher temperatures), as well as pH 5.5–6, 7 and 9. As shown in **Fig. 3**, no other phosphorylating agent was as effective as AcP at synthesising ATP in the presence of Fe^3+^. The only other phosphorylating agent to show any notable efficacy was carbamoyl phosphate (CP), which is similar in structure to AcP; it has a carbamate (-CO-NH_2_) rather than acetate (-CO-CH_3_) bound to phosphate. CP produced about half the ATP yield of AcP at 20 °C and pH 5.5–6 (**Fig. 3a**), but barely a quarter of the yield at pH 7 (**Fig. 3b**). At pH 9, only cyclic trimetaphosphate (cTMP) produced any ATP at all, albeit after a delay of more than 20 hours (**Fig. 3c**).

**Figure 3.**
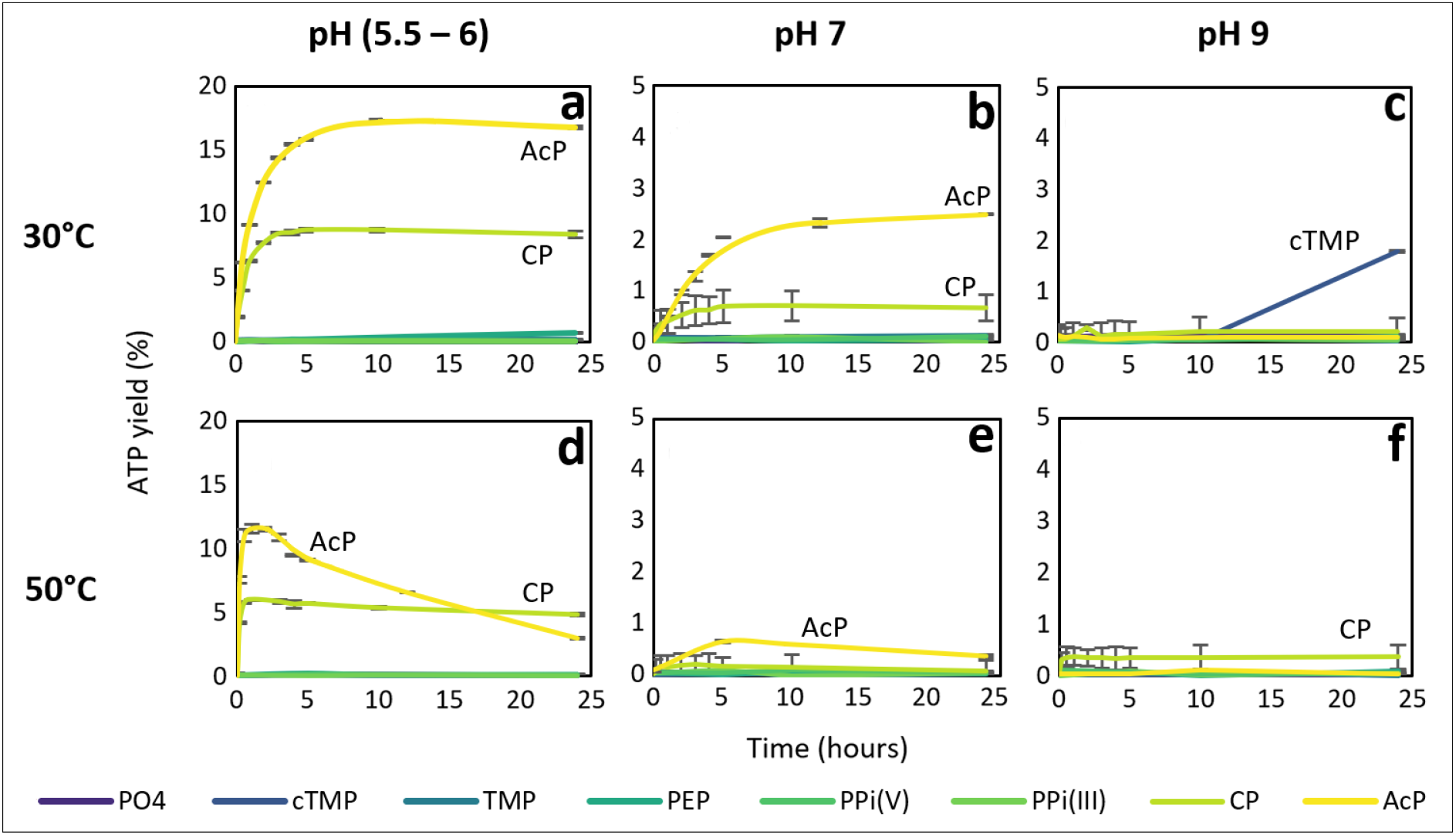
ATP synthesis with different phosphorylating agents. 1:4 ADP:phosphorylating agent reaction catalysed by Fe^3+^ with various phosphorylating agents at different pH and temperature. PO_4_: potassium phosphate; cTMP: trimetaphosphate; TMP: trimethyl phosphate; PEP: phosphoenolpyruvate; PPi(V): pyrophosphate; PPi(III): pyrophosphite; CP: carbamoyl phosphate; AcP: acetyl phosphate. N = 3 ±SD.

At 50 °C, CP generated ATP continuously over 24 hours at pH 5.5–6, despite producing only half the yield in the first 2 hours. The fact that ATP yield declined over time with AcP indicates that ATP was hydrolysed over hours at 50 °C; it was not replenished because AcP also hydrolysed at that temperature [10]. While CP has a similarly low thermal stability, the primary decomposition product is cyanate [57], which is itself a proficient condensing agent [58]. This likely contributed to a balance between the synthesis and hydrolysis of ATP over 24 hours. Only AcP formed any ATP at 50 °C and pH 7 (**Fig. 3e**), consistent with the pH sensitivity of CP seen at 30 °C. CP did form ATP at low yield at 50 °C and pH 9 (**Fig. 3f**), and we can infer again that it is due to the decomposition product cyanate;. The main conclusion here is that from a panel of eight plausibly prebiotic phosphorylating agents, only AcP was capable of generating an ATP yield of > 10% in water at both 30 and 50 °C. The only other agent to show remotely comparable efficacy at mildly acidic pH was CP, but its maximal yield was half that of AcP. The fact that CP was capable of synthesising ATP at low yield under warm alkaline condition (50 °C, pH 9) in fact lowers its phosphorylating potential as it is less capable of sustaining a disequilibrium of ATP/ADP ratio in a dynamic pH environment (see Discussion).

### Phosphorylation of ADP to ATP is unique among nucleotide diphosphates

We next explored the propensity of AcP to phosphorylate other canonical nucleotide diphosphates (NDPs), specifically cytidine diphosphate (CDP), guanosine diphosphate (GDP), uridine diphosphate (UDP) and inosine diphosphate (IDP). While not a canonical base, inosine is the precursor to both adenosine and guanosine in purine synthesis. Importantly, from a mechanistic point of view, inosine lacks the amino group incorporated at different positions onto the purine rings of adenosine and guanosine, but like GDP, IDP has an oxygen in place of the N6 amino group of adenosine. The results clearly show that AcP will phosphorylate ADP but not other NDPs (**Fig. 4a-e**), demonstrating a strong dependence on the structure of the nucleobase. For all NDPs, a peak for the corresponding triphosphate was present at the start of the reaction, but this did not change over 3 hours for any NDP except ADP. As noted above for ADP, the presence of the NTP at 0 h can be ascribed to minor contamination of the commercial standard during the manufacturing process.

**Figure 4.**
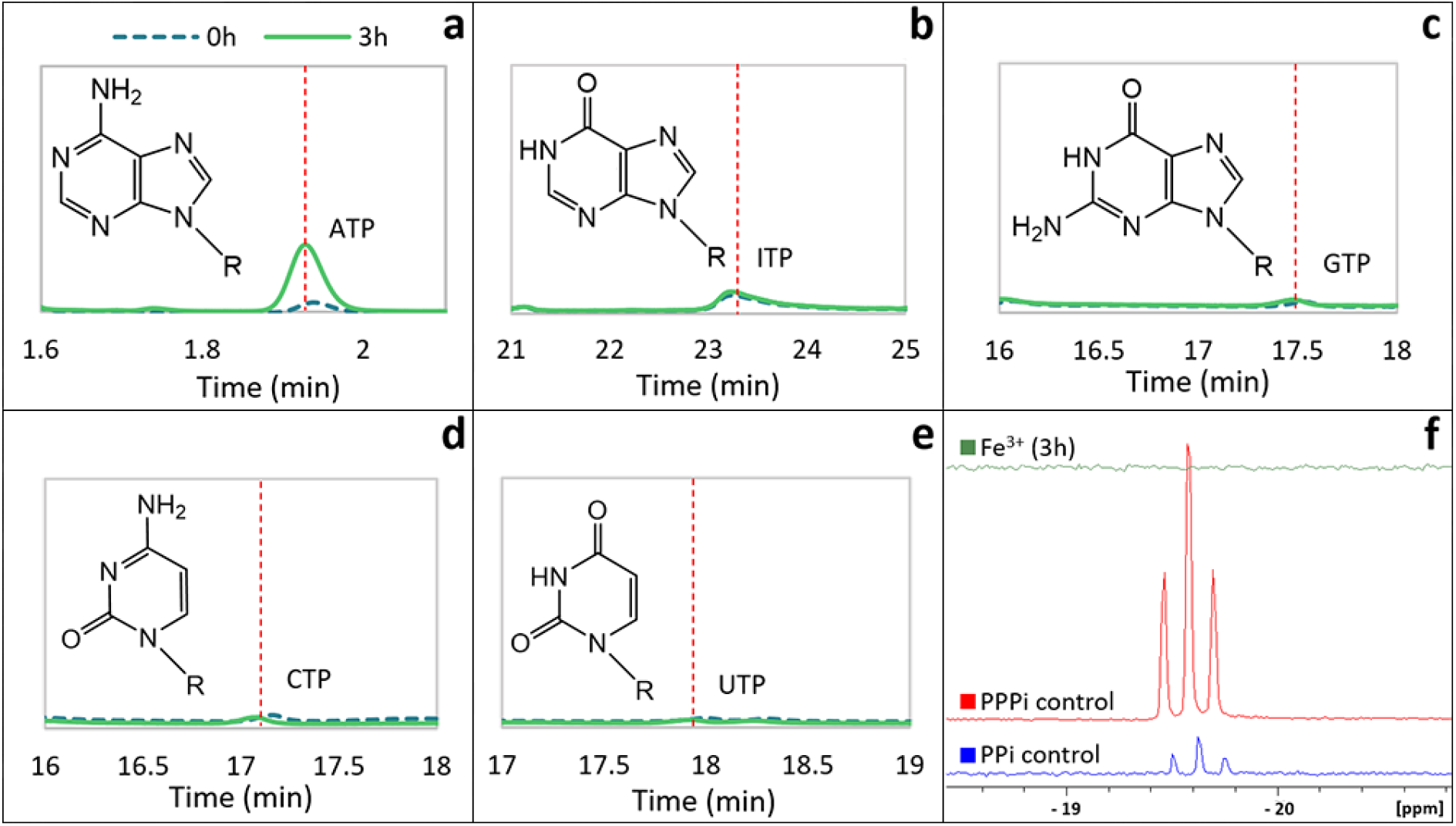
Phosphorylation of nucleotide diphosphates by AcP. HPLC chromatogram of the resulting NTP of the phosphorylation of (**a**) adenosine diphosphate (ADP), (**b**) inosine diphosphate (IDP), (**c**) guanosine diphosphate (GDP), (**d**) cytidine diphosphate (CDP) and (**e**) uridine diphosphate (UDP) by AcP catalysed by Fe^3+^ at 30°C and pH ∼5.5–6 at the beginning of the reaction (0 h, broken line, blue) and after 3 hours (solid line, red). The molecular structure of each base forming the nucleotides is shown. (**f**) ^31^P–NMR spectrum of PPi (bottom, blue), PPPi (middle, red) and the reaction PPi (1 mM) + AcP (4 mM) + Fe^3+^ (500 μM) at 30°C and pH ∼5.5–6 after 3h (top, green).

To explore the dependence of phosphorylation on the nucleobase, and to establish whether Fe^3+^ interacts directly with the base as well as its diphosphate tail, ADP was substituted by potassium pyrophosphate (PPi) in the reaction mixture with AcP and Fe^3+^. No triphosphate was detected by ^31^P-NMR (**Fig. 4f**), which suggests that the adenine ring does indeed need to interact directly with Fe^3+^. We note that Fe^3+^ heavily interferes with ^31^P–NMR spectroscopy due to its paramagnetism. To minimize the presence of Fe^3+^ in the sample, we therefore performed solid-phase extraction twice before NMR. Despite this precaution, the experimental samples still showed some deformation, suggesting that Fe^3+^ continued to interact with the phosphate groups (**SI Fig. 4**) [59]. Nonetheless, this small deformation is cosmetic and does not conceal the absence of triphosphate in the reaction mixture. We also considered whether Fe^3+^ could interact with the adenine ring but not the diphosphate tail, analysing the phosphorylation of AMP to ADP. AcP did indeed phosphorylate AMP to ADP in the presence of Fe^3+^ (**SI Fig. 5**) but at considerably lower yield than ADP to ATP. Thus, Fe^3+^ interacts preferentially with the purine ring coupled to the diphosphate tail.

The fact that neither pyrimidine NDP could be phosphorylated suggests that the purine ring (or at least adenosine) is essential for positioning the interactions between Fe^3+^ and AcP. ADP has an amino group at N6, whereas GDP has a carbonyl at C6 and an amino group at N2; inosine has a carbonyl group at C6; and both GDP and IDP have a protonated N at N1. We infer that the critical moiety in the adenosine ring for phosphorylation by AcP with Fe^3+^ as catalyst must be the N6-amino group of adenosine, as the IDP and GDP ring structures are equivalent elsewhere. In particular, from a mechanistic point of view, we note that the N7 is equivalent in all three purine rings, so although this might also interact with Fe^3+^, as suggested by others [60–63], it cannot be the critical moiety.

### Catalysis of ADP phosphorylation does not involve nucleotide stacking

To understand how Fe^3+^ catalyses the phosphorylation of ADP to ATP, we tested the effect of varying the Fe^3+^ ion concentration. Holding the ADP and AcP concentrations constant at 1 mM and 4 mM, respectively, we varied the Fe^3+^ concentration from 0.05 to 2 mM. We found that the maximal ATP yield was produced by 1 mM Fe^3+^, indicating that the optimal ADP:Fe^3+^ stoichiometry of the reaction was 1:1 (**Fig. 5a**). Following Kitani *et al*. [33]we confirmed that low concentrations of either Mg^2+^ or Ca^2+^ (up to 2 mM) slightly increased the ATP yield in the presence of 1 mM Fe^3+^. This suggests that either of these divalent cations can stabilise the newly formed ATP and liberate Fe^3+^ to catalyse the next phosphorylation of ADP (**Fig. 5a**).

**Figure 5.**
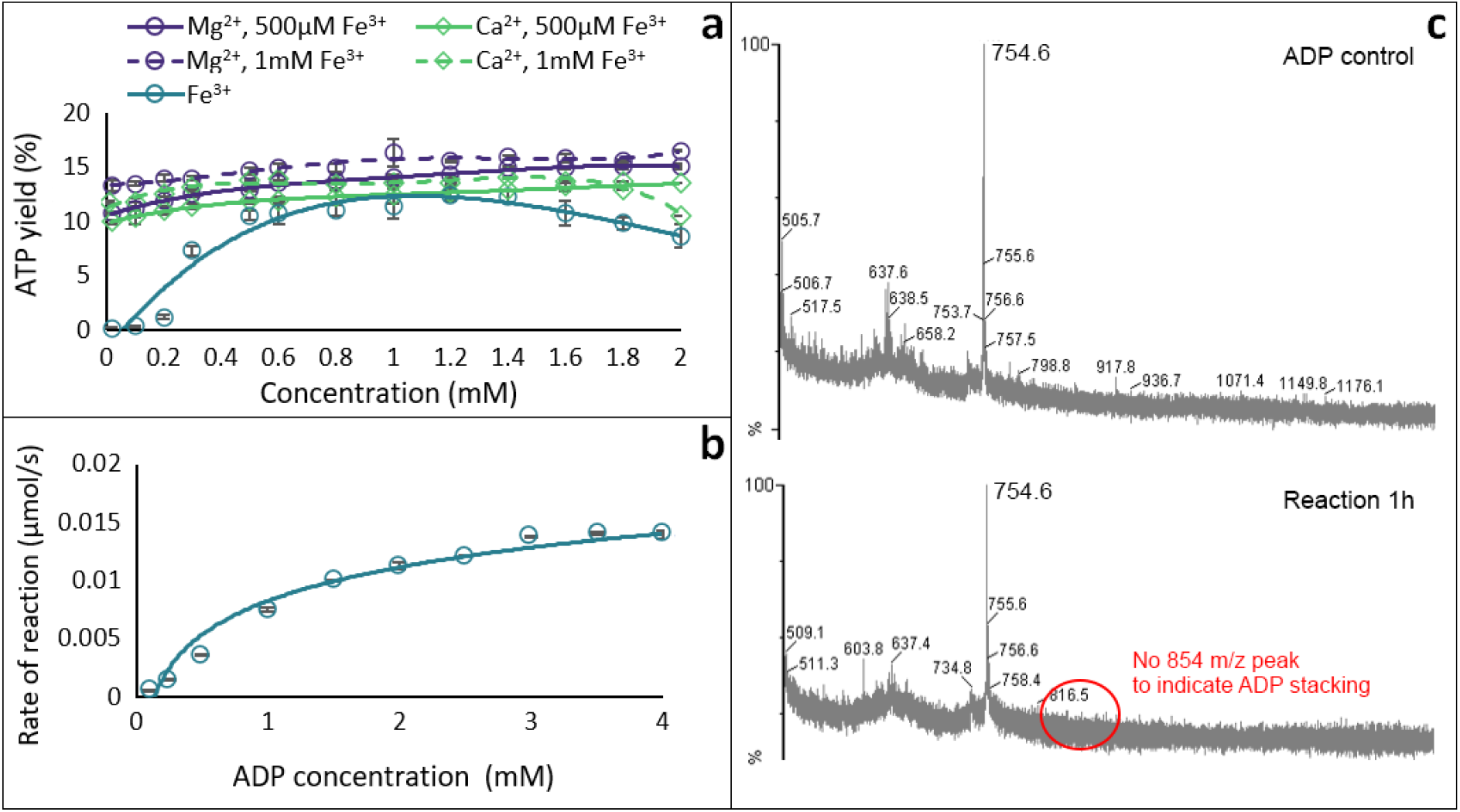
Mechanism studies. (a) Effect of varying concentration of Fe^3+^ (teal circles) and adding increasing concentrations of Mg^2+^ (purple circles) and Ca^2+^ (green diamonds) on ATP yield at 2 h from the reaction ADP (1 mM) + AcP (4 mM) with 0.5 mM Fe^3+^ (solid line) and 1mM Fe^3+^ (broken line) at 30°C and pH ∼5.5–6. (N = 3 ±SD and 2 ±SD, respectively). (b) Michaelis-Menten kinetic analysis on the ADP + AcP reaction catalysed by Fe^3+^ (0.5 mM). N = 3 ±SD. (c) MALDI-ToF spectra of ADP control (top) and a reaction sample at 1 h (bottom).

We next conducted a kinetic study of the phosphorylation reaction, specifically varying the ADP concentration and monitoring the reaction rate. The resulting curve resembled a characteristic Michaelis-Menten mechanism for an enzyme, indicating that Fe^3+^ does indeed act as a catalyst (**Fig. 5b**). The question remained whether a single Fe^3+^ was interacting directly with a single ADP and AcP, or whether larger units such as stacked ADP rings were involved. Stacking can alter the geometry of which group interacts with Fe^3+^ (**SI Fig. 6**) and has previously been suggested as a possible mechanism[64]. However, MALDI-ToF analysis, which can sensitively detect stacked nucleotides, showed no difference between the ADP control and the reaction sample; the main visible peaks appeared to be dimers of ADP/AMP present in the commercial ADP standard, possibly due to freeze-drying during production of ADP [65] (**Fig. 5c**). This demonstrates that stacking of ADP to coordinate the Fe^3+^ ion does not occur as a mechanistic step in the reaction. That in turn constrains more tightly which groups in the base could potentially interact with metal ions such as Fe^3+^.

Altogether, our results suggest that the high charge density of Fe^3+^ allows it to interact directly with the N6 amino group on the adenine ring, while anchoring AcP in position for its phosphate group to interact with the diphosphate tail of ADP, giving a taut conformation of ADP (**Fig. 6a**). The interaction with the dianion has been proposed before [66,67] and is key because at the optimal pH of 5.5–6, the first two hydroxyl groups of ADP (p*K*_a_ 0.9 and 2.8) are deprotonated, while the external OH group (p*K*_a_ 6.8) remains protonated, and is therefore not available for nucleophilic attack [68]. The interaction of the two deprotonated OH groups with Fe^3+^ has the effect of lowering the p*K*_a_ of the outermost OH group, thus deprotonating it and enhancing its nucleophilicity (**Fig. 6b**). The phosphate group of AcP is readily positioned for nucleophilic attack by the newly deprotonated O^−^ of ADP, forming ATP (**Fig. 6c**). This mechanism also explains why Ca^2+^ and Mg^2+^ slightly increase the rate of reaction; these ions are able to displace Fe^3+^ from the ATP product (as they interact better with the triphosphate tail; **Fig. 6d**), freeing the Fe^3+^ to catalyse further reactions (**Fig. 6e**).

**Figure 6.**
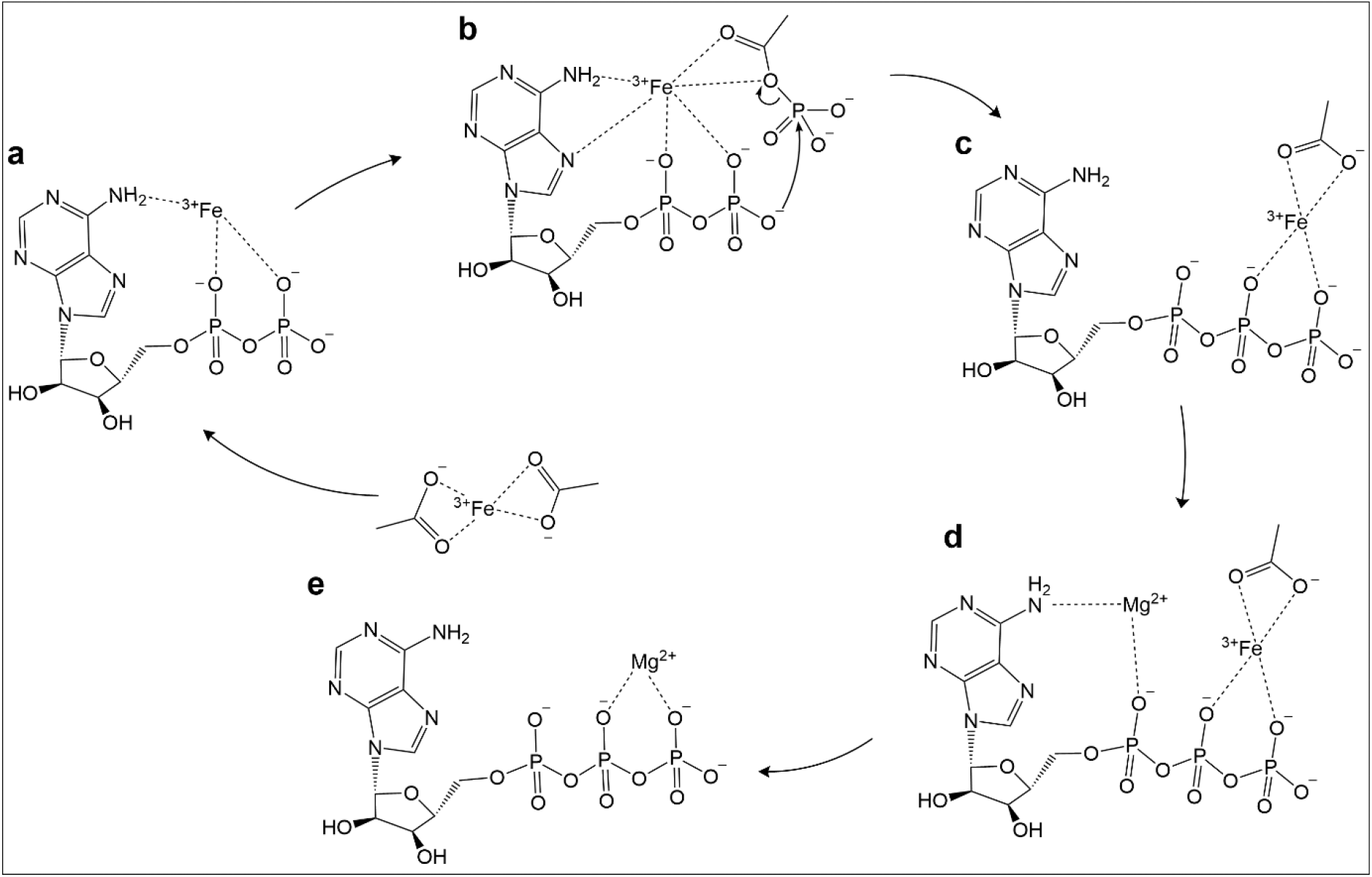
Potential mechanism. Fe^3+^, stabilised by the 6-NH_2_ and N7 groups on adenine, interacts with the dianion of ADP, lowering the p*K*_a_ of the outermost OH group, enhancing nucleophilicity. Fe^3+^ interacts with the oxygens of a molecule of the surrounding AcP, bringing it close enough to facilitate the phosphate transfer. Fe^3+^ then moves from Pα to the Pβ and Pγ of ATP and ultimately abandons the ATP chelated by acetate groups facilitated by the favourable association of Mg^2+^. Fe^3+^ is then available to catalyse another phosphorylation of ADP.

## Discussion

Our results support the following conclusions: (i) acetyl phosphate (AcP) efficiently phosphorylates ADP to ATP, but only in the presence of Fe^3+^ ions as catalyst (**Fig. 1**); (ii) the reaction takes place in water and can occur in a wide range of aqueous environments (**Fig. 2**); (iii) no other phosphorylating agent tested was as effective as AcP (**Fig. 3**); and (iv) adenine is unique among canonical nucleobases in facilitating the phosphorylation of its nucleotide diphosphate to the triphosphate (**Fig. 4)**. Taken together, these findings suggest that the pre-eminence of ATP in biology has its roots in aqueous prebiotic chemistry. The substrate-level phosphorylation of ADP to ATP by AcP is uniquely facilitated in water under prebiotic conditions and remains the fulcrum between thioester and phosphate metabolism in bacteria and archaea today [2]. This implies that ATP became the universal energy currency of life not as the endpoint of genetic selection or some frozen accident, but for fundamental chemical reasons, and probably in a monomer world before the polymerization of RNA, DNA and proteins.

The work presented here provides a compelling basis for each of these statements, but also raises a number of questions. Why ferric iron? Unlike AcP or ATP itself there is no clear link with biology in this case; we had expected other ions more commonly associated with nucleotides, notably Mg^2+^ or Ca^2+^ [39,40], to play a more clear-cut role. In fact, their catalytic effect was only noticeable in the presence of Fe^3+^, as has been reported before, whereas higher concentrations, equivalent to modern ocean conditions, precluded ATP synthesis. We infer that the reason Fe^3+^ plays a unique role relates in part to its high charge density and small ionic radius. The fact that only ADP could be phosphorylated among canonical nucleobases suggests that Fe^3+^ interacts directly with the N6 amino group on the adenine ring as well as the N7 previously noted by others [60–63]. But the interactions between Fe^3+^ and the N7 moiety alone cannot explain our results, as no triphosphate was formed in the absence of the N6-amino group, for example in the case of GDP. The fact that ADP is phosphorylated more readily than AMP (**SI Fig. 5**) indicates that Fe^3+^ also interacts with the diphosphate tail of ADP. And the fact that the optimal stoichiometry of Fe^3+^ to ADP is 1:1, coupled with the absence of evidence for stacking of bases by MALDI-ToF (**Fig. 5**), indicates that a single Fe^3+^ ion interacts with a single ADP, and necessarily also with a single AcP.

As shown in **Fig. 6**, these stipulations require an unusually taut molecular configuration of ADP, far from the loose conformation usually depicted, if only for ease of presentation. The orientation of the adenine ring in relation to metal ions has long been disputed, with some arguing that it should face the opposite way in apposition to the phosphate tail [69]. Others have suggested an equivalent orientation to that proposed here [67,70], some specifically with Fe^3+^ [60,61]. In any case, this taut conformation almost certainly requires the interacting ion to have a high charge density and small ionic radius, to draw each of these groups into close enough proximity to react. Among the cations tested here, Fe^3+^ has the highest charge density and the smallest ionic radius [71]. Nonetheless, some of the other ions studied, notably Cr^3+^ and Co^3+^, have a similar ionic radius and charge density, yet do not have a remotely comparable catalytic effect, so the size and charge density cannot be the only explanation for our results. The electronic configuration of Fe^3+^ may also play a role: unlike Cr^3+^ and Co^3+^, Fe^3+^ has the electronic configuration [Ar]3d^5^, having all 5 d orbitals half occupied. However, Mn^2+^, which can substitute Mg^2+^ in the catalytic centre of acetate kinase, has an equivalent 3d orbital, yet yielded negative results in our experiments. If so, then size, charge density and electronic configuration might all play a role. These possibilities need to be explored in future work.

Why acetyl phosphate? The idea that this small (2-carbon) molecule might have acted as a phosphoryl donor at the origin of life has a long history, going back to Lipmann himself [10,13,17,19,24–29], as indeed does its confounding potential as an acetyl donor. Acetyl phosphate still plays a global signalling and energy transduction role in bacteria [72], in part because its free energy of hydrolysis (and therefore its phosphorylating potential) is greater than that of ATP (ΔG°′ = -43 kJ mol^-1^ versus -31 kJ mol^-1^, respectively). When complexed in a 1:1 ratio with ADP, therefore, AcP has the potential to transfer its phosphate to form ATP, and so serves as a labile energy source in cells, linked to the excretion of acetate as waste. But the actual change in ΔG depends on how far from equilibrium the ratio of AcP/Ac + Pi or ATP/ADP + Pi has been pushed, and hence varies depending on conditions. In our experiments, all phosphoryl donors were added at equivalent excess. The fact that the ΔG°′ for hydrolysis of PEP (−62 kJ mol^-1^) and CP (−51 kJ mol^-1^) are markedly greater than that for AcP means that free-energy change is only part of the explanation for the efficacy of AcP. The fact that ATP was primarily formed by AcP in the presence of Fe^3+^ ions instead implies that the critical factors were (i) the position of the two phosphoester oxygen atoms in relation to the Fe^3+^, and (ii) the phosphate group in relation to the diphosphate tail of ADP, as shown in **Fig. 6**. In other words, both AcP and ADP are favoured not for selective or thermodynamic reasons, but kinetic – because their chemistry is facilitated by molecular geometry in aqueous prebiotic environments.

The only other molecule with equivalent geometry in this regard is carbamoyl phosphate (CP), which our model would therefore predict should have some phosphorylating efficacy. CP was indeed the only other species to show significant phosphorylating activity in our system (**Fig. 3**). CP has long been considered as a plausible prebiotic phosphorylating agent [55,73–75] and can also promote the formation of ATP in the presence of Ca^2+^ or Ba^2+^ ions [55,76–78]. Like AcP, CP retains a place in modern metabolism, for example as a substrate for carbamate kinase, phosphorylating ADP to ATP in microbial fermentation of arginine, agmatine, and oxalurate/allantoin [79], as well as the *de novo* synthesis of pyrimidines (although not as a phosphorylator) [80]. Taken together with our own results, these findings suggest that both AcP and CP are molecular ‘living fossils’ of prebiotic chemistry, retaining a role in modern metabolism due to their felicitous chemistry. But despite these similarities, CP was less effective than AcP at generating ATP under mildly acidic to neutral conditions (**Fig. 3**). This difference holds important connotations for its ability to power work.

A major question for prebiotic chemistry is how can an energy currency power work? As noted in the Introduction, there is nothing special about the bonds in ATP; rather, the ATP synthase powers a disequilibrium in the ratio of ADP to ATP, which amounts to 10 orders of magnitude from equilibrium in the cytosol of modern cells. Only that disequilibrium powers work; no equilibrium mixture of ATP and ADP can power anything. But molecular engines such as the ATP synthase use ratchet-like mechanical mechanisms to convert environmental redox disequilibria into the highly skewed ratio of ADP to ATP [81]. How could a simple prebiotic system composed of monomers sustain a disequilibrium in the ratio of ATP to ADP that powers work? One possibility is that the environment itself could sustain critical disequilibria across short distances, such as membranes. The fundamental disequilibrium that drives work in essentially all cells is the proton-motive force – at its simplest, the difference in proton concentration, or pH, across membranes. This mechanism is highly relevant to ATP, given the strong dependence of ATP synthesis versus hydrolysis on pH, specifically because the phosphorylation potential of ATP depends on its free energy of hydrolysis, which increases with pH [82,83]. Far from being an environmental limitation, the narrow pH range facilitating ATP synthesis reported here may therefore help to drive work in a monomer prebiotic world.

Dynamic environments such as alkaline hydrothermal systems can sustain steep pH gradients across thin inorganic barriers, as mildly acidic Hadean ocean waters (pH 5-6) continually mix with strongly alkaline hydrothermal fluids (pH 9-11) in microporous labyrinths that operate as electrochemical flow reactors [28,84–86]. We have previously shown that thin inorganic barriers containing FeS minerals such as mackinawite can sustain proton gradients as steep as 4 pH units across single 25 nm FeS nanocrystals [87]. Such steep pH gradients could in principle operate across protocells as well as inorganic barriers. Alkaline hydrothermal conditions promote the self-assembly of protocells with bilayer membranes composed of mixed amphiphiles (fatty acids and fatty alcohols) [88]). These protocells can bind to mineral surfaces, potentially exposing them to the steep pH gradients across barriers [89]. Equivalent pH gradients can drive the synthesis of organics including formate [90] and potentially thioesters [31]. The critical point is that proton flux across membranes in hydrothermal systems could promote the phosphorylation of ADP to ATP under locally acidic conditions close to the barriers, followed by hydrolysis linked to phosphorylation under more alkaline conditions in the cytosol of protocells. At face value, the ATP yield reported here at pH 5-5–6 after 10 hours was 17.4 % (corresponding to 156.5 µM) while the yield at pH 9 was 0.043 %, corresponding to 0.4 µM, a difference of 400-fold. Thus, a geologically sustained difference in pH across membranes could drive a local disequilibrium in the ATP/ADP ratio of 2-3 orders of magnitude, enough to power work even in the absence of other possible factors such as temperature. Higher temperatures (50 °C) promote both the rapid synthesis and hydrolysis of ATP (**Fig. 2b**), which should amplify this driving force. We stress that these considerations require further elucidation, but in principle steep pH gradients can drive a disequilibrium in the ATP/ADP ratio that powers work.

Are these far-from-equilibrium conditions consistent with the high water-activity and low ion requirements for optimal ATP synthesis in our experiments? High concentrations of Mg^2+^ (50 mM) and Ca^2+^ (10 mM) precluded ATP synthesis, implying that this chemistry would not be favoured in modern oceans, but would be feasible in freshwater systems. Likewise, ferrous iron could be oxidized to ferric iron by photochemical reactions or oxidants such as NO derived from volcanic emissions, meteorite impacts or lightning strikes, which also points to terrestrial geothermal systems as a plausible environment for aqueous ATP synthesis [91]. But it is less clear if steep gradients (of pH or anything else) could sustain disequilibria in ATP/ADP ratios in terrestrial geothermal systems. In any case, our results certainly do not rule out the alkaline hydrothermal systems discussed above. Some shallow submarine systems such as Strytan in Iceland are sustained by meteoritic water, and feature Na^+^ gradients as well as H^+^ gradients [92]; such mixed systems could have been common in shallow Hadean oceans. The concentration of divalent cations in the Hadean oceans may also have been lower than modern oceans, with estimates varying widely [40,93]. Regardless of mean ocean concentrations, strongly alkaline conditions tend to precipitate Ca^2+^ and Mg^2+^ ions as aragonite and brucite, so their concentration can be much lower in hydrothermal systems.

Ferric iron may also have been available, even in deep ferruginous oceans. Thermodynamic modelling shows that the simple mixing of alkaline hydrothermal fluids with seawater in submarine systems can promote continuous cycling between ferrous and ferric iron, potentially forming soluble hydrous ferric chlorides [94] (which our experiments show to have the same effect as ferric sulphate, **SI Fig. 7**). The availability of ferric iron is critical for other prebiotic catalysts including cysteine-FeS clusters [38,95–97] and has been discussed in more detail elsewhere [38]. Other conditions considered here, including salinity and pressure [44], have only limited effect on ATP synthesis in warm alkaline hydrothermal systems. Finally, we do not envision ATP synthesis taking place in open geochemical systems, but rather within leaky protocells composed of mixed amphiphiles [88] and capable of simple metabolic heredity [98]. Any system that can generate nucleotides will likely also form carboxylic acids such as citrate that can chelate divalent cations. We therefore consider our results to be consistent with a wide range of prebiotic aqueous environments.

This is not the first report of ATP synthesis at moderate yield under prebiotic conditions. What do our findings add to earlier work? We here provide an unexpected link between aqueous prebiotic chemistry and biochemistry. For example, earlier work using cyanate as a condensing agent generated ATP from ADP [99,100], but cyanate also phosphorylated other nucleotide diphosphates. That was important as it showed that biologically relevant condensations are possible in water, but differed from modern biochemistry in that cyanate does not feature in extant metabolism, nor does it discriminate between bases. Cyanate therefore gives little insight into the origins of biochemistry as we know it, and specifically, the question of why ATP is the universal energy currency.

The work reported here shows that AcP is unique among a panel of relevant phosphorylating agents in that it can phosphorylate ADP to ATP in the presence of Fe^3+^. AcP is formed readily through prebiotic chemistry, and remains central to prokaryotic metabolism, making it the most plausible precursor to ATP as a biochemical phosphorylator [10]. Critically important, AcP does not phosphorylate other nucleotide diphosphates, giving a compelling insight into how ATP came to be so dominant in modern metabolism. Our findings indicate that the high charge density and electronic configuration of Fe^3+^ can position molecules in water to react in the absence of macromolecular catalysts such as RNA or proteins, or even mineral surfaces. Beyond that, our results suggest that steep pH gradients could in principle generate disequilibria in the ratio of ATP to ADP of several orders of magnitude, enabling ATP to drive work even in a prebiotic monomer world. Once formed, ATP would promote intermediary metabolism through phosphorylation and as a precursor to cofactors, notably NADH, FADH and coenzyme A, while also driving the polymerization of amino acids and nucleotides to form RNA, DNA and proteins, via liberation of pyrophosphate as the leaving group. If so, then ATP became established as the universal energy currency for reasons of prebiotic chemistry, in a monomer world before the emergence of genetically encoded macromolecular engines.

## Materials and Methods

### Materials

All salts were purchased from Sigma-Aldrich, except for copper nitrate hemipentahydrate (Cu(NO_3_)_2_·2.5H_2_O), copper sulphate pentahydrate (CuSO_4_·5H_2_O) and manganese nitrate hexahydrate (Mn(NO_3_)_2_·6H_2_O, Alfa Aesar), TEAA (triethylammonium acetate, Fluka), and CTP (Cytidine 5’- triphosphate sodium salt, Cambridge Bioscience). All solvents were HPLC-grade and purchased from Fischer. All reagents used were analytical grade (≥ 96%).

### Reaction setup

Depending on the solubility of the analytes, reactions were carried out in either a stationary (SciQuip HP120-S) or a shaking (ThermoMix HM100-Pro) dry block heater.

For the reaction, stock solutions of di-nucleotides (sodium salts, ≥96%, Sigma-Aldrich), phosphorylating agents and metal catalyst were freshly prepared as to avoid freeze-thawing (10 mM for reactions to be analysed via HPLC, 1 M for reactions to be analysed via NMR). Except where indicated, the ratios of analytes in a solution were 1(ADP):4(AcP) and 1(Fe^3+^):2(ADP). When needed the pH was adjusted using aqueous HCL and NaOH (1 M or 3 M)

After checking the pH (Fisher Scientific accumet AE150 meter with VWR semi-micro pH electrode), samples were taken at time-points (0, 10 and 30 min, 1 to 5, 10 and 24h) and, unless otherwise specified, immediately frozen at −80 °C for next-day analysis.

#### Pressure reactor

Experiments under pressure were performed in a pressure vessel (Series 4600-1L-VGR with single inlet valve, Parr Instrument Company), pressurised with N_2_ gas and placed on a hotplate (Fisherbrand Isotemp Digital Stirring Hotplate) at 30°C. Samples for both the high pressure experiment and ambient pressure control experiment were prepared in 2 mL glass headspace vials (Agilent Technologies) whose caps were pierced with a needle.

#### FeS clusters

FeS clusters coordinated by 5mM of L-cysteine were prepared under anaerobic conditions and water sparged with N_2_ was used to prepare all solutions. Stock solutions of 10 mM Na_2_S, 10 mM FeCl_3_, 50 mM of L-cysteine and 1 M of NaOH were prepared either in water or in 10 mM bicarbonate buffer (pH 9.1). A volume of 4 mL of Na_2_S and 4 mL of L-cys were added to 28 mL of water/buffer, and the pH adjusted to ∼9.8 using NaOH. A volume of 4 mL of FeCl_3_ was then added and the volume adjusted to 40 mL to obtain a 1 mM FeS solution.

Oxygen levels in the anaerobic glovebox were maintained below 5 ppm when possible, and no work was conducted if this level was surpassed.

#### UV/Vis Spectroscopy

UV/Vis spectroscopy was used to verify the formation of FeS clusters. A volume of 1 mL of FeS stock solution was placed in a crystal cuvette, which was sealed with parafilm under anaerobic conditions. Spectra were obtained using a Thermofisher NanoDrop 2000c, with a baseline correction of 800 nm.

### Analysis

#### HPLC

Samples were prepared at collection by spinning at 4,000rpm for 2 minutes and diluting 200 µL in 800 µL of EDTA solution (500 µL in 100 mM PO_4_ buffer at pH 7.1) prior to freezing, in order to chelate the Fe^3+^ ions in solution that would otherwise block the HPLC column.

Thawed samples were filtered using syringe filters (ANP1322, 0.22 μm PTFE Syringe filter, Gilson Scientific Ltd.) attached to a 1 mL sterile syringe (BD Plastipak Syringes) in 2 mL headspace vials and analysed on an HPLC instrument (Agilent Technologies, 1260 Infinity II); peaks were identified using pure standards. The wavelengths for UV detection were usually set at 254 nm and 260 nm (most suitable for cyclic rings such as adenosine), while the column tray temperature was maintained at room temperature. Two different columns were used depending on the pH of the sample being analysed: Poroshell 120 EC-C18 for pH 2-8 and Poroshell HPH-C18 for pH 9-11.

Mobile phase A consisted of 80 mM phosphate buffer (made by mixing equal parts of potassium phosphate dibasic (40 mM) and potassium phosphate monobasic (40 mM) salts dissolved in water) adjusted to pH 5.8 using 3 M HCl and filtered with 0.2 μm nylon membrane filters (GNWP04700, 0.2 μm pore size, Merck Millipore Ltd.), while mobile phase B consisted of 100% methanol. The injection volume was 1 μL, with a flow rate of 1 mL/min, and the run was an isocratic gradient that consisted of 95% B for 5 minutes.

For experiments using nucleotide diphosphates with different bases, analyses were carried out on a Polaris C18-A column, with mobile phase A consisting of 10 mM potassium phosphate monobasic buffer with 10 mM Tetrabutylammonium hydroxide (TBAH) adjusted to pH 8 using 3 M HCl and filtered with 0.2 μm nylon membrane filters (GNWP04700, 0.2 μm pore size, Merck Millipore Ltd.), while mobile phase B consisted of 100% methanol (method described in Table 2). The wavelengths for UV detection were set at 254, 260, and 271 nm for guanosine, uridine and inosine, and cytidine, respectively.

**Table 2.** HPLC method for G, C, I and U nucleotides experiments.

|  |  |
| --- | --- |
| **Mobile phase A** | 10 mM KH<sub>2</sub>PO<sub>4</sub> + 10 mM TBAH in HPLC-grade water |
| **Mobile phase B** | 100% HPLC-grade methanol |
| **Gradient** | 5% B → 50% B (up during 25 min) → 50% B (for 2 min) → 95% B (up during 6 sec) → 95% B (for 3 min) → 5% B (down during 6 sec) → 5% B (for 2 min) |
| **Flow rate** | 1.5 mL/min |
| **Injection volume** | 1 µL |

Two flush methods (Table ***3*)** were employed to preserve the column: Flush 1 was used every 12-15 samples, then three rounds of Flush 1 followed by one run of Flush 2 were run prior to switching off the machine.

**Table 3.** HPLC flush methods. These are run at the end of a set number of sample analyses computational analysis was done using Agilent OpenLAB software (ChemStation Edition). Each peak was manually integrated using the calibration curves as reference and the raw file was exported for data manipulation. As residual ATP is present in the ADP commercial standard, the yield of the reaction is calculated by subtracting the reading for ATP at timepoint 0 from all subsequent timepoint readings.

|  | *FLUSH 1* | *FLUSH 2* |
| --- | --- | --- |
| **Mobile phase A** | HPLC-grade water | HPLC-grade water |
| **Mobile phase B** | 100% HPLC-grade methanol | 100% HPLC-grade methanol |
| **Gradient** | 5% → 95% B (up during 15 minutes) → 95% B (for 5 min) → 5% B (down during 10 min) | Initial: 5% B (for 17 min) → 95% B (up during 18 min) → 95% B (for 17 min) → 60% B (down during 6 min) → 60% B (for 17 min) |
| **Flow rate** | 1 mL/min | 1 mL/min |

#### ^31^P–NMR

As iron is paramagnetic and thus tampers with NMR spectra, samples prepared for ^31^P-NMR were purified using solid phase extraction (SPE) after thawing. The SPE cartridge (InertSep ME-1, 300mg/3mL) was equilibrated with 3mL of 100% methanol and then washed with 3mL H_2_O, after which the sample was passed through and collected. The procedure was tested on control samples to ensure appropriate recovery

A volume of 0.9 mL of purified sample was added to 0.1 mL of D_2_O and dispensed in an NMR tube (Norell Standard Series 5mm Precision NMR Sampling Tubes) for analysis (^1^H decoupling, Bruker Avance 400 MHz, 52 scans). The data was processed using the Bruker TopSpin 4.0.7 software and peaks were identified using pure standards.

#### ESI MS

Electrospray Ionisation Mass Spectrometry was used to confirm the identity of ATP through MS/MS. After purification through SPE (see previous section) the sample was loaded into a 0.5 mL glass syringe (Gastight Syringe Model 1750 RN, Hamilton) and directly infused into the mass spectrometer (Finnigan LTQ Linear Ion Trap mass spectrometer) at a flow rate of 10 μL/min. To avoid contaminations, the syringe and line were flushed with 100% methanol before and after sample infusion, and the spectra recorded.

The mass spectrometer was operated in negative ion mode and the capillary voltage was set at -16 V. Data were collected from 100 to 2000 *m/z* with an acquisition rate of 5 spectra per second. For the MS/MS Ar was used as the collision gas and the collision energy was adjusted to 30 eV. The software Xcalibur (Thermo Scientific) was used for method setup and data processing.

#### MALDI-ToF MS

Samples were thawed and desalted using a protocol adapted from Burcar *et al*.[101]. Two solvents were prepared: an ACN solution consisting of 50% acetonitrile in water and a 0.1 M TEAA solution in water.

Using a Millipore C18 zip tip (Sigma), 10 µL of ACN solution were aspirated and discarded 3 times. The three rinses were repeated with 10 µL of the TEAA solution. To allow for the retention of the analyte by the zip tip matrix, 10 µL of sample were aspirated up and down eight times and then discarded. A volume of 10 µL of water were aspirated and discarded, followed by 10 µL of the TEAA solution and once again 10 µL of water. A volume of 4 µL of ACN were slowly aspirated up and down three times and deposited into a small Eppendorf microcentrifuge tube.

The MALDI-ToF protocol used was designed by Whicher *et al*. [10]. The matrix consisted of 2,4,6-trihydroxyacetophenone monohydrate (THAP) and ammonium citrate dibasic, and was freshly prepared before the analysis using equal volumes of stocks that were maintained at 4°C for a maximum of a week.

A volume of 2 μL of matrix solution was mixed with 2 μL of sample, deposited onto a clean steel MALDI-ToF plate and allowed to evaporate for 30 minutes before the introduction of the steel plate into the instrument (Waters micro MX mass spectrometer). The analytical conditions were: reflectron and negative ion mode, 400 au of laser power, 2000 V of pulse, 2500 V of the detector, 12,000 V of flight tube, 5200 V of reflector, 3738 V of negative anode, and 500– 5000 amu of scan range. The mass spectrometer was calibrated using a low-molecular-weight oligonucleotide standard (comprising of a DNA 4-mer, 5-mer, 7-mer, 9-mer, and 11-mer (Bruker Daltonics)). Each oligonucleotide standard was initially dissolved in 100 μL water, divided in aliquots and frozen at −80 °C. A fresh aliquot was used at each analytical calibration.

## Pinna *et al*. Supplementary Information

**SI Figure 1.**
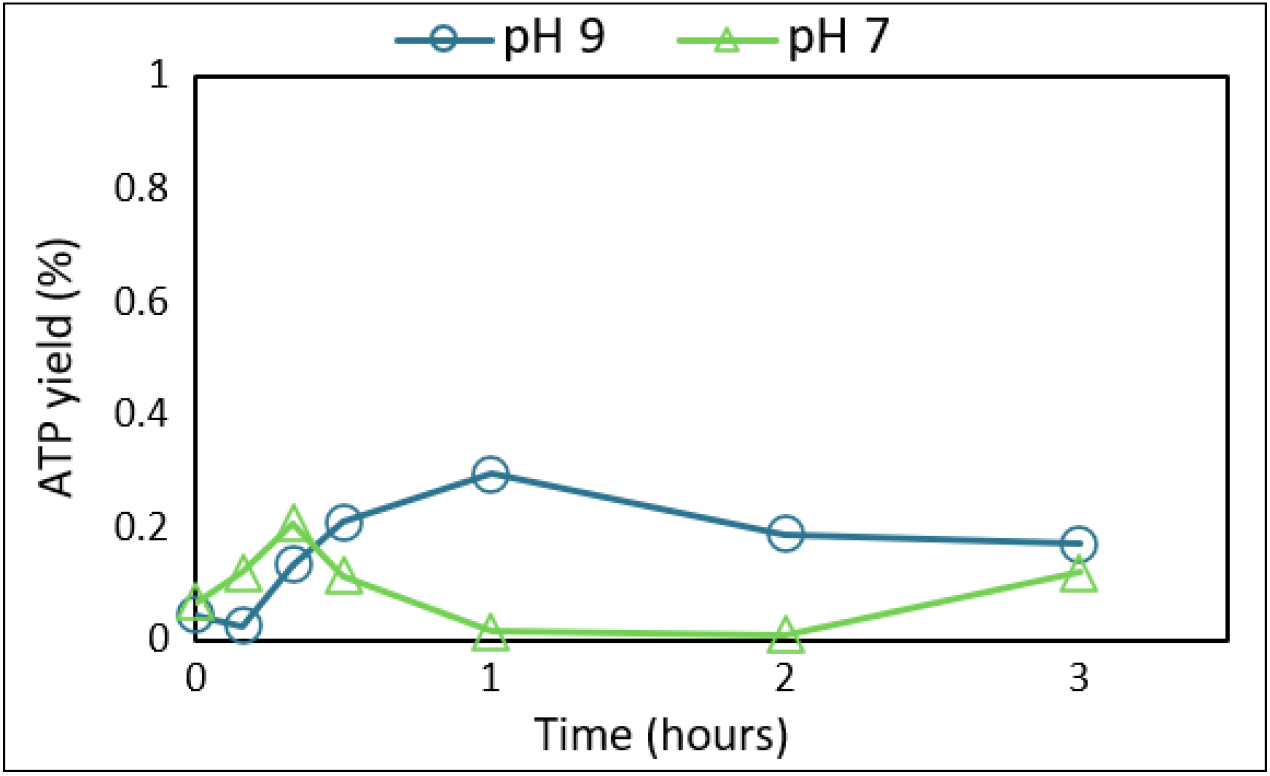
Comparison of the ATP yield from the reaction ADP (1 mM) + AcP (4 mM) at 30°C in a FeS clusters-rich 10 mM bicarbonate solution at pH 9 (circles, teal) and 7 (triangles, green).

**SI Figure 2.**
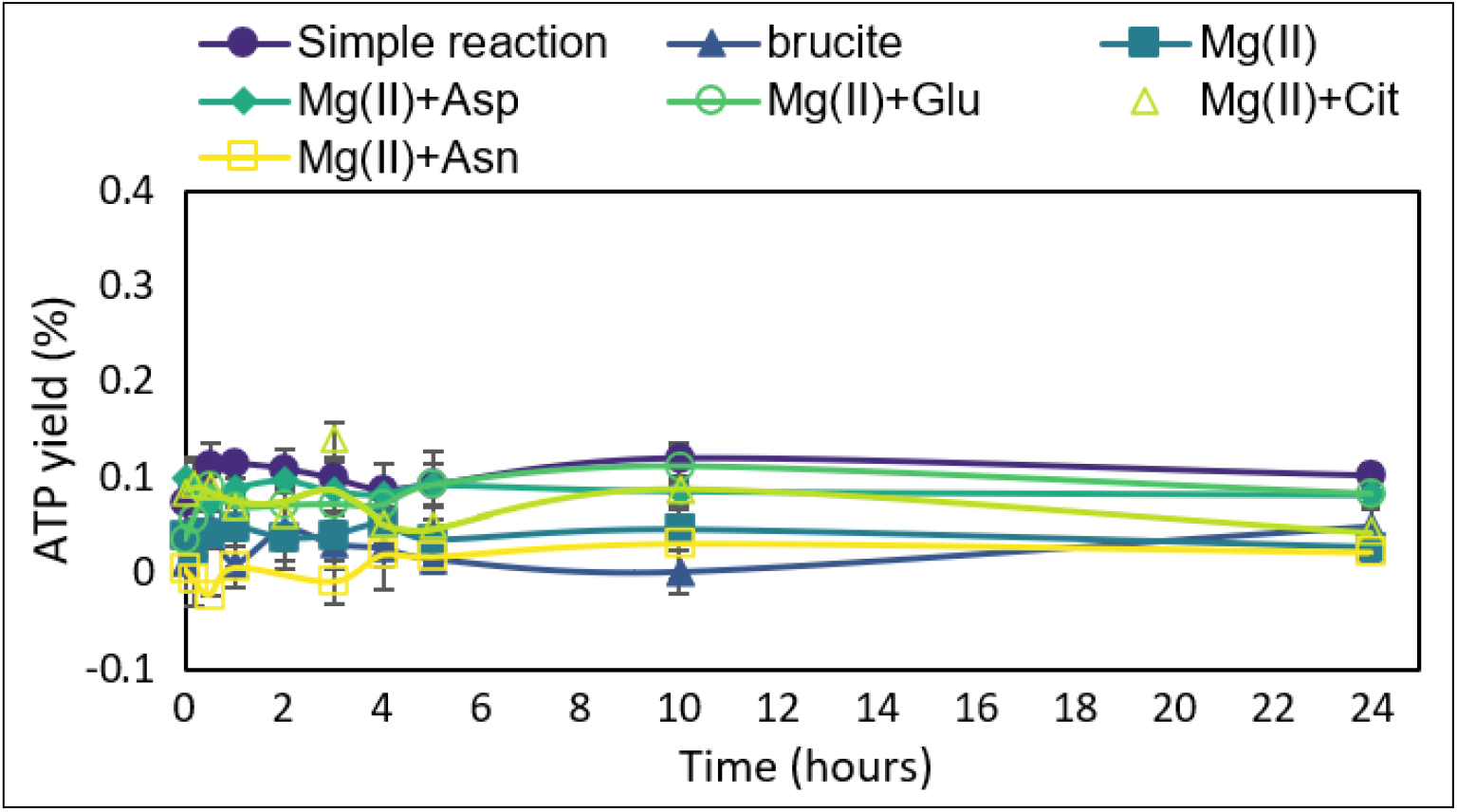
Comparison of reaction ADP (1 mM) + AcP (4 mM) at 30°C and pH ∼5.5–6 with different forms of magnesium (ionic form and mineral form brucite) coordinated by citrate or amino acids. N = 3 ±SD. Brucite is a hydroxide mineral (Mg(OH)_2_) with a unit structure reminiscent of the Mg^2+^ coordination by aspartate in enzymes such as Mg^2+^-dependent RNA polymerase.

**SI Figure 3.**
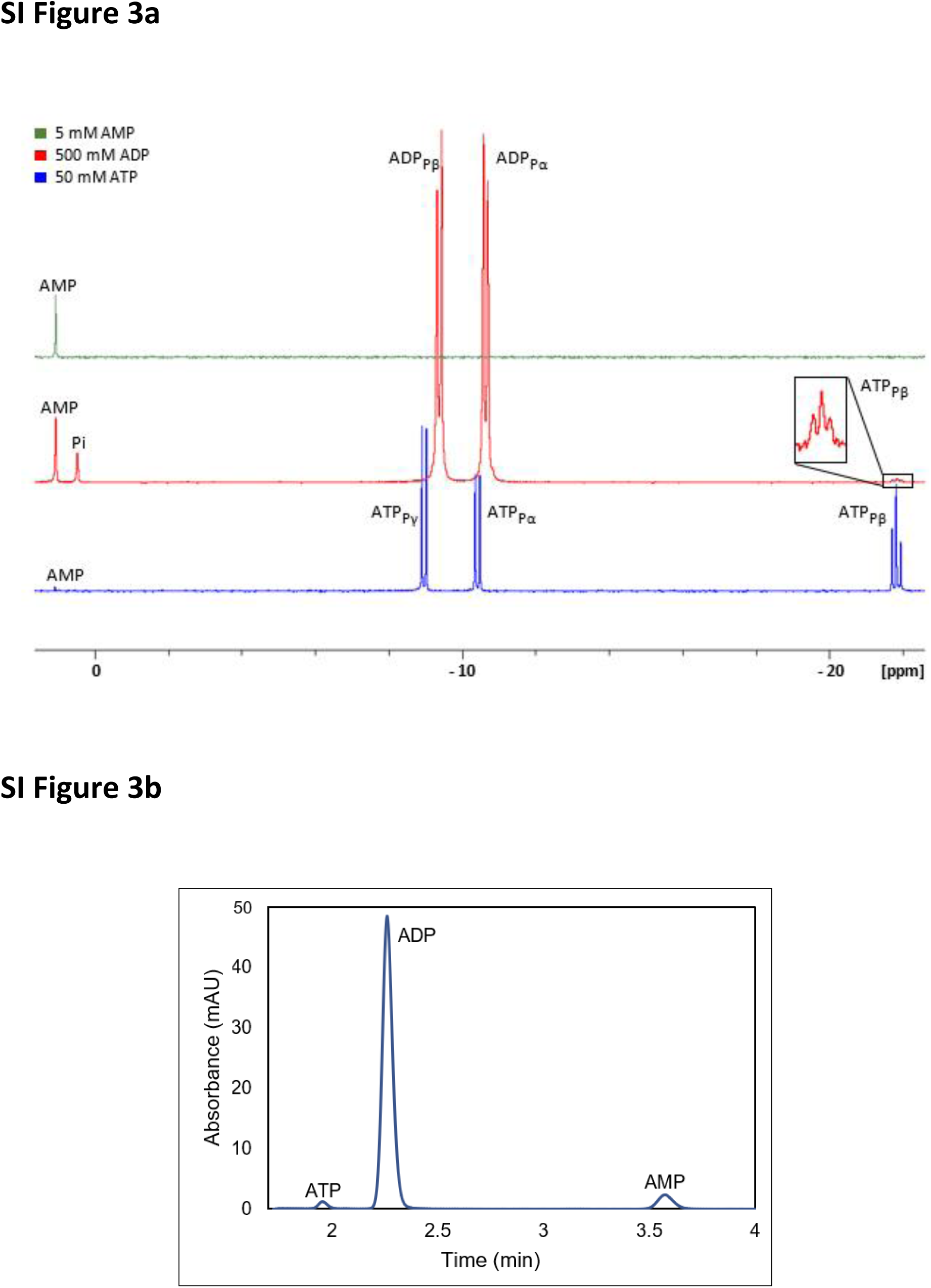
Residual levels of ATP present in the ADP commercial standard. (a) Comparison of ^31^P-NMR spectra of commercial AMP (green), commercial ADP (red), and commercial ATP (blue); the graph insert shows a zoomed in area of ATP signal. (b) HPLC chromatogram of commercial ADP (1mM). All peaks labelled for clarity.

**SI Figure 4.**
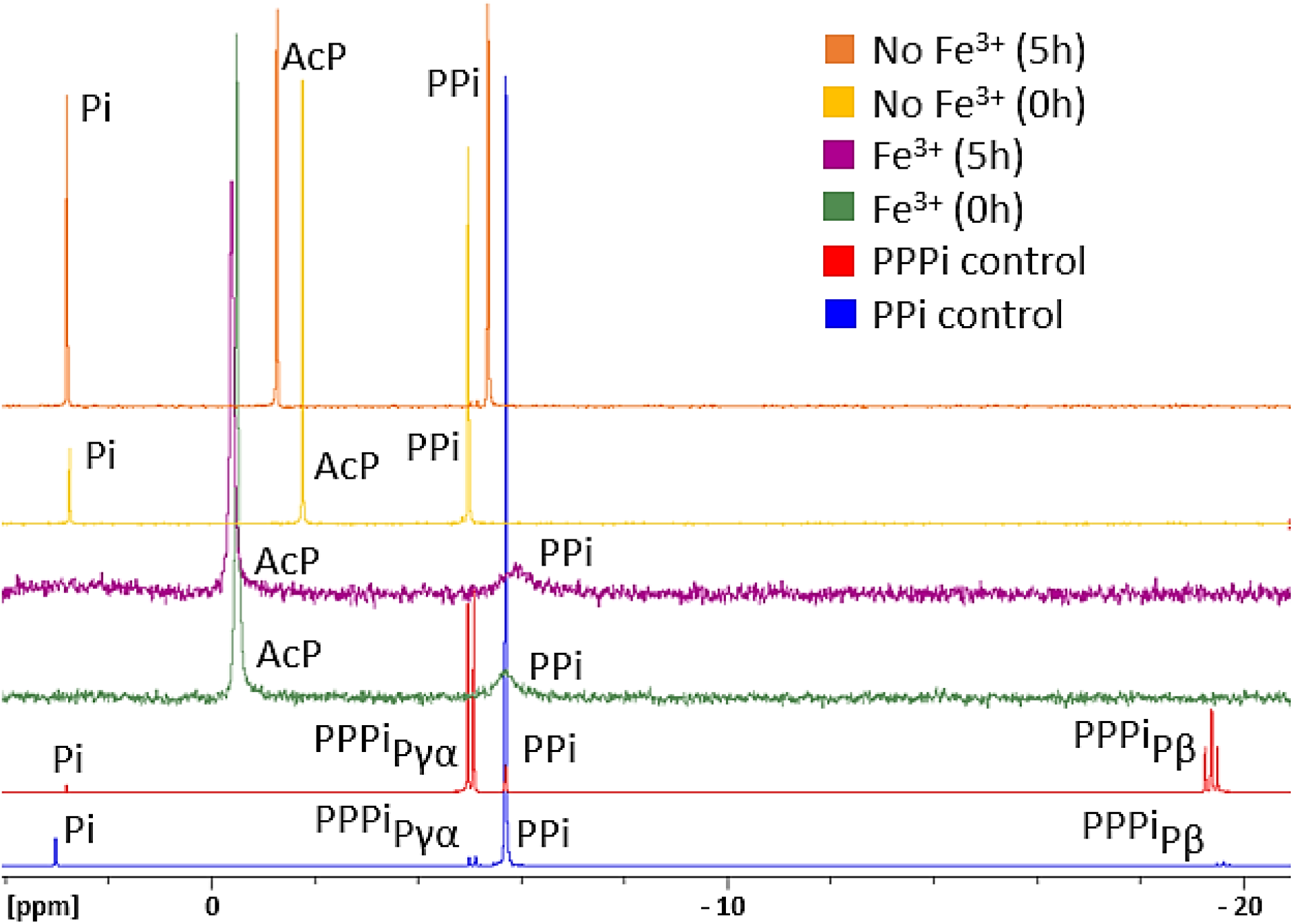
Comparison of ^31^P-NMR spectra of the phosphorylation of PPi by AcP in the absence (orange and yellow) and presence (purple and green) of Fe^3+^, commercial PPPi (red), and commercial PPi (blue). All peaks labelled for clarity.

**SI Figure 5.**
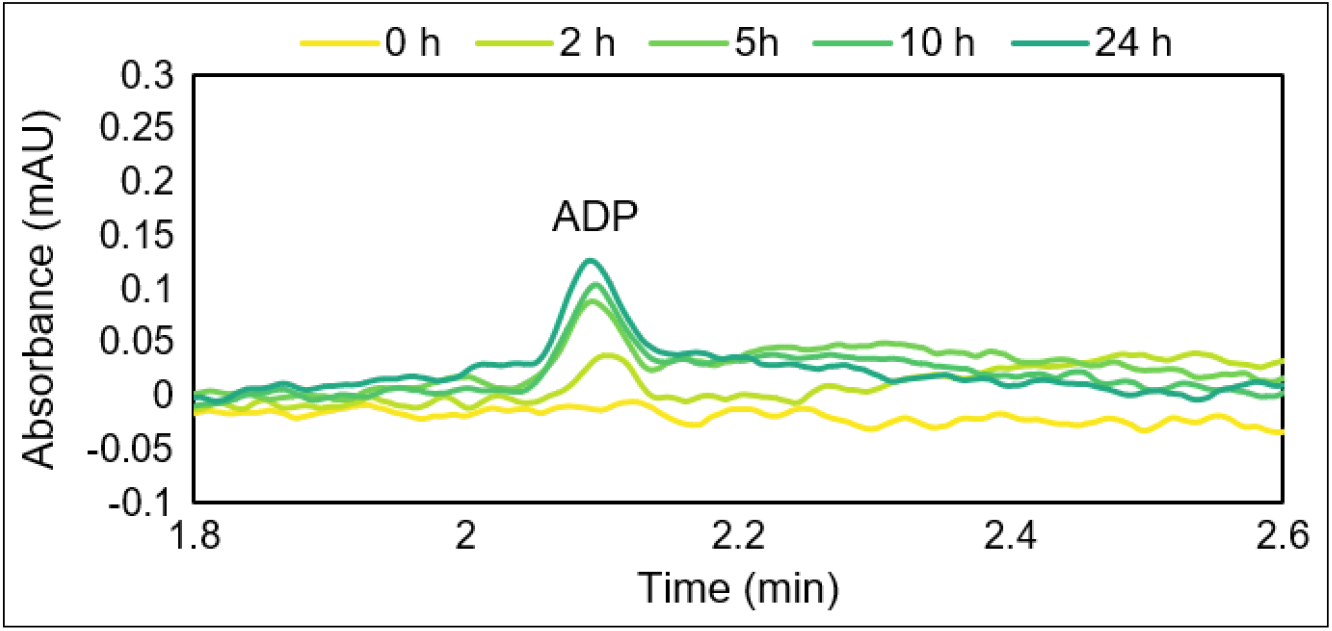
HPLC chromatogram showing the progressive ADP synthesis over 24 hours via phosphorylation of AMP by AcP in the presence of Fe^3+^ at 30°C.

**SI Figure 6.**
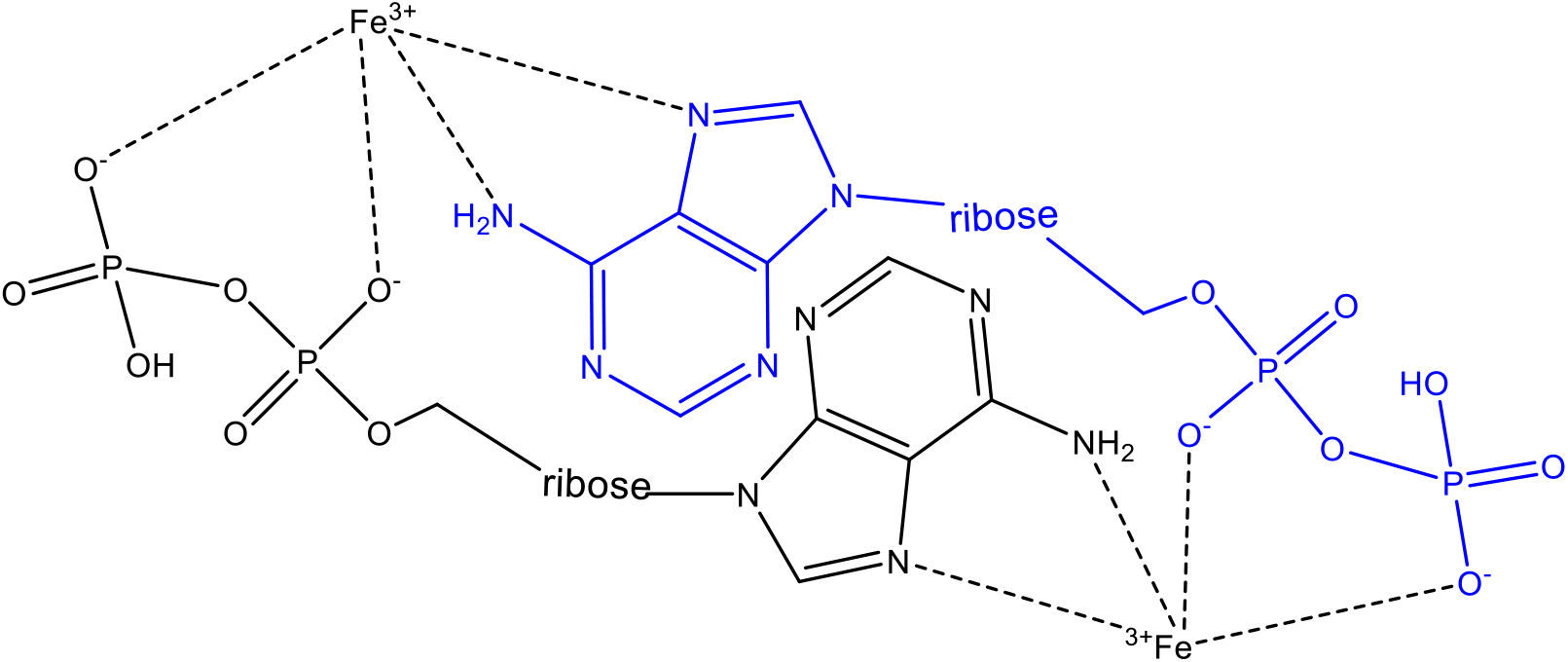
Possible stacking of ADP coordinated by Fe^3+^.

**SI Figure 7.**
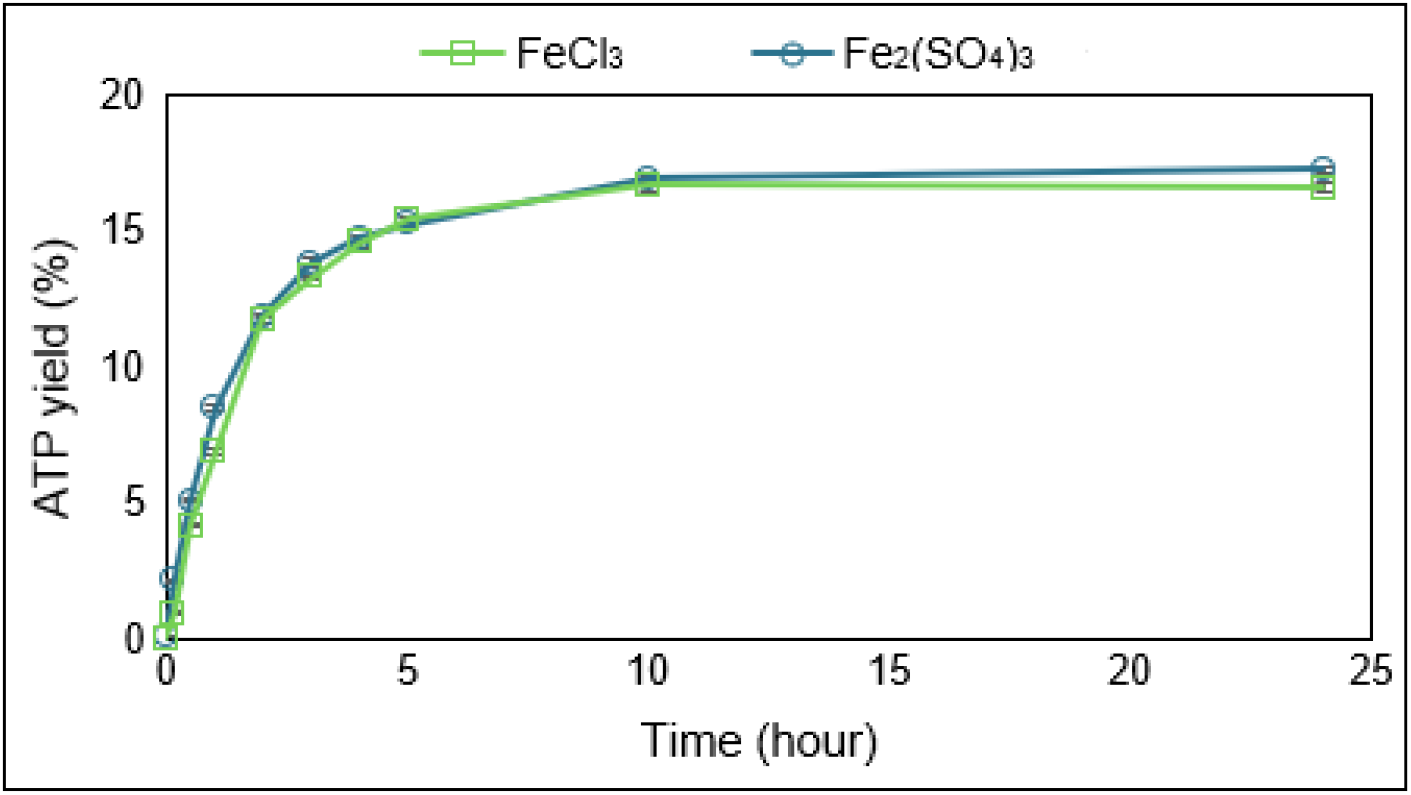
Comparison of the ATP yield from the reaction ADP (1 mM) + AcP (4 mM) + Fe^3+^ (0.5 mM) at 30°C where the Fe^3+^ is given by either FeCl_3_ (squares, green) or Fe_2_(SO_4_)_3_ (circles, teal). N = 3 ±SD

## Notes

### Competing Interest Statement

The authors have declared no competing interest.

